# Hypothalamic CYP46A1 cholesterol metabolism regulates diet-induced obesity

**DOI:** 10.64898/2026.01.10.698773

**Authors:** David V.C. Brito, Adriana Arrulo Pereira, Renata Sandres de Souza Araujo, Bruno André Martins Várzea, Xavier Guerreiro Anastácio, Telma Cristina Aureliano Castro, Célia Aveleira, Sara Carmo-Silva, Marisa Ferreira-Marques, Ana Luísa Sousa-Coelho, José Matos, Leonor Faleiro, Nathalie Cartier, Sandro Alves, Clévio Nóbrega

## Abstract

Obesity is a growing global health challenge which affects over 38% of the population and increases the risk of developing metabolic disorders. Effective long-term treatment options remain limited and often lead to significant side effects. The hypothalamus plays a central role in metabolic regulation by integrating peripheral and central signals to maintain energy homeostasis. While the hypothalamus regulates peripheral lipid metabolism, the role of lipid metabolic pathways within the hypothalamus itself remains unclear. Therefore, the molecular mechanisms linking hypothalamic activity to the development of obesity remain poorly understood. Given that cholesterol is the most abundant brain lipid, we investigated molecular players involved in its local regulation. Here, we identify the rate-limiting enzyme for brain cholesterol degradation, Cyp46a1, as a critical regulator of diet-induced obesity. We show that Cyp46a1 expression is significantly reduced in the hypothalamus of C57BL/6J mice exposed to a high-fat diet, suggesting a role in metabolic dysregulation. We used recombinant Adeno-Associated Viruses (AAV) to modulate Cyp46a1 expression in the arcuate nucleus, a key hypothalamic region regulating energy balance. We evaluated longitudinal changes in body weight, adipocyte size, PPAR-γ expression, and hepatic lipid accumulation and pancreatic histopathology in mice undergoing chow or high fat diets. These experiments demonstrated that hypothalamic Cyp46a1 regulates body weight, adipocyte size, and hepatic and pancreatic histopathology. Conversely, CYP46A1 overexpression protects against high-fat diet-induced obesity and improves glucose homeostasis and insulin sensitivity, likely via crosstalk with hepatic and pancreatic function. Moreover, we found that hypothalamic CYP46A1 expression selectively modulates gut microbiota composition, linking this enzyme to the microbiota-gut-brain axis. Finally, we found that Cyp46a1 levels influence cognitive and motor performance, suggesting broader physiological relevance. Collectively, our findings reveal a previously unrecognized role of hypothalamic Cyp46a1 and cholesterol pathway control in obesity pathophysiology and metabolic homeostasis. These findings identify hypothalamic Cyp46a1 as a promising therapeutic target for obesity and its related comorbidities.

## Introduction

About 38% of the global population is overweight or obese and this number is predicted to rise to 51% by 2035^1^. Obesity increases the risk of developing debilitating disorders such as diabetes *mellitus* type 2, cardiovascular diseases and cancer, thereby contributing to a higher global mortality rate^2^. Current therapeutic options are limited, relying primarily on lifestyle changes and invasive interventions such as bariatric surgery or treatment with glucagon-like peptide-1 (GLP-1) agonists, which are often associated with adverse effects^3^. Therefore, understanding the pathophysiological processes underlying obesity is essential for developing innovative and more effective therapeutic strategies.

The hypothalamus plays a central role in regulating energy homeostasis by integrating peripheral and central signals to maintain metabolic balance. Dysregulation of hypothalamic function has been implicated in various forms of obesity^4^. For instance, hypothalamic obesity can arise from lesions caused by conditions such as tumor growth^5^. Moreover, the hypothalamus also contributes to cholesterol metabolism via the autonomic nervous system^6–8^, which regulates peripheral responses to high caloric intake. Cholesterol turnover in the central nervous system (CNS) is mainly regulated by the cholesterol 24-hydroxylase (CYP46A1) enzyme, which converts cholesterol into 24S-hydroxycholesterol, a product that can cross the blood-brain-barrier (BBB) for systemic elimination. While the hypothalamus regulates peripheral lipid metabolism, the role of lipid metabolic pathways within the hypothalamus itself remains unclear. Therefore, the molecular mechanisms linking hypothalamic activity to the development of obesity remain poorly understood. We and others have shown that CYP46A1 levels are decreased in various neurodegenerative disorders^9^, making it a promising therapeutic target^9^. Nonetheless, a possible role in the metabolic regulation of obesity remains largely unexplored.

In this study, we investigated whether CYP46A1 in the hypothalamus plays a role in diet-induced obesity. We found that mRNA levels of this enzyme are decreased in the hypothalamus of C57BL/6J mice exposed to a high-fat diet. To investigate its functional significance, we modulated CYP46A1 expression using recombinant adeno-associated viruses (rAAVs) to either knock-down or overexpress this enzyme in the arcuate nucleus (ARC), a key hypothalamic region involved in regulating energy intake and expenditure^10^. We then assessed its impact on food intake, body weight, white adipose tissue (WAT) morphology, and systemic metabolic parameters under chow and high-fat diet (HFD) conditions.

We further investigated the downstream effects of CYP46A1 modulation on key obesity-associated features, including hepatic steatosis, insulin sensitivity, and gut microbiome composition. Our findings revealed that hypothalamic CYP46A1 levels regulate body weight and adipocyte size, while also modulating organism-wide metabolic and behavioral changes associated with diet-induced obesity. We further demonstrate that hypothalamic CYP46A1 contributes to obesity-induced impairments in glucose homeostasis and insulin sensitivity, likely mediating these effects through systemic interactions with hepatic and pancreatic functions. Finally, our results suggest that hypothalamic CYP46A1 influences gut microbiota composition as well as cognitive and motor behaviors, offering novel insights into the multifaceted role of this enzyme in obesity.

## Materials and Methods

### Mice

Male and female C57BL/6J mice (Charles River, Spain) were used for this study. Mice were group-housed (4 mice per cage) on a 12h light/dark cycle with, 22 ± 2°C and 55 ± 15% relative humidity. Animals were randomly assigned to two dietary groups, with the overall ratio of male to female mice balanced. One group had access to a low-fat control diet (Chow, D12450J, 10% fat, Research Diets, USA) or to a high-fat diet (HFD, D12492, 60% fat, Research Diets, USA). Chow and HFD diets as well as autoclaved water were provided *ad libitum*. Mice were accommodated to the diets for four weeks prior to stereotaxic surgery. All behavioral experiments took place during the light phase except open field test which was also conducted during dark phase. Sick and/or injured mice from cage-mate fighting were excluded from this study. The experiments were carried out in accordance with the European Community Council directive (2010/63/EU) for the care and use of laboratory animals. Researchers received adequate training (FELASA) and certification to perform the experiments from Portuguese authorities (*Direção-Geral de Alimentação e Veterinária*).

### Recombinant Adeno Associated Virus (rAAV) production

The AAV vectors were produced and purified by Atlantic Gene therapies (Inserm U1089, Nantes, France). The viral construct AAV5-shCyp46a1 contained the expression cassette of a short hairpin RNA (shRNA) targeting the mouse *Cyp46a1* gene, driven by an U6 promoter and a *GFP* reporter gene driven by the phosphoglycerate kinase 1 (PGK1) promoter in the adeno-associated viral vector serotype 5 (AAV5)^10^.The AAV5 vectors encoding shCyp46a1 sequences were generated by transient transfection of human embryonic kidney (HEK) 293T cells and purified on caesium chloride ultracentrifugation gradients, as previously described^11^. The viral construct for AAVrh10-CYP46A1 contained an expression cassette consisting of cDNA encoding the human CYP46A1 protein, driven by a cytomegalovirus (CMV) early enhancer/chicken β-actin synthetic promoter (CAG) and surrounded by inverted terminal repeat (ITR) sequences of AAV. GFP reporter gene driven by the phosphoglycerate kinase 1(PGK1) promoter in the adeno-associated viral vector serotype 5 (AAV5) as previously described ^12^.

### Body weight, food and water consumption monitoring

Food and water amounts were registered every two weeks. This data was used to calculate food and water intake per g/ml of body weight (BW). The ratio of food ingested (g) per total BW for cage (g) was calculated and then multiplied for the body weight of each mouse to obtain the food intake [(food ingested/total BW for cage) X BW of each mouse)]. The ratio of water ingested (mL) per total body weight for cage (g) was calculated and then multiplied for mouse BW to obtain the water intake per week.

### Open-Field test

Mice underwent stereotaxic delivery of rAAVs into the arcuate nucleus of the hypothalamus and 4 and 8-weeks later were subjected to the open field test for the assessment of locomotor and anxiety-like behavior^13,14^. Briefly, mice were placed in an acrylic square box (40×40×40cm), in a sound-attenuating controlled room, illuminated with a white light during the tests. The bottom of the box was divided into nine equally sized squares to define the center zone (center square, about 11% of total area) and the remaining eight squares were defined as the periphery zone. The animals were gently placed in the box and their movement activity was recorded during 10 min in daytime period and 5 min in the nighttime period using a GoPro Hero camera 42 (GoPro Inc., USA). The videos were automatically analyzed using the ANY-Maze behavioral tracking software (Stoelting Company, Europe).

### Y Maze spontaneous alternation test

Mice underwent stereotaxic delivery of rAAVs into the arcuate nucleus of the hypothalamus and 4 and 8-weeks later were subjected to the Y Maze test as previously described^15^. Briefly mice were placed at the beginning of one arm and their movement were recorded for 5 minutes, to evaluate the total number of arm entries, total distance travelled and spontaneous alternations. Each animal is expected to show a tendency to alternate between arms, instead of going back to the previously visited. Videos were automatically analyzed using ANY-Maze behavioral tracking software (Stoelting Company, Europe). The percentage of spontaneous alternations was calculated according to the following formula:

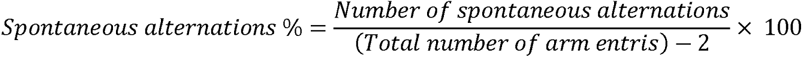

### Sample preparation and protein extraction

Mice were anesthetized with halothane followed by decapitation. Hypothalamic tissue was rapidly microdissected and frozen until extraction. QIAzol (Qiagen) was added and later manually homogenized with a syringe and needle. D-chloroform was added to the homogenate and mixed for 15 seconds. The homogenate was left for 3 minutes at room temperature and then centrifuged at 12000g for 15 minutes at 4°C. The aqueous phase was collected for RNA extraction and 100% ethanol was added to the interphase and phenol phases. The samples were mixed carefully and incubated at room temperature (RT) for about 3 minutes. After, the samples were centrifugated at 2000g for 2 minutes at 4°C. To the supernatant (protein fraction) was added isopropanol to allow protein precipitation and was mixed for 15 seconds. The total volume was incubated at room temperature for 10 minutes. Later, the samples were centrifuged at 12000g for 10 minutes at 4°C and the supernatant was removed. Guanidine-ethanol solution was added to and let to incubate at RT for 20 minutes. After, the samples were centrifugated at 7500g for 5 minutes at RT and the supernatant was removed. The addition of guanidine-ethanol, incubation, centrifugation, and removal of supernatant steps were repeated 3 times. 100% Ethanol was added to the samples, vortexed and incubated at room temperature for 20 minutes. After, the samples were centrifuged at 7500g for 5 minutes at RT, the supernatant was removed, and the pellet air dried for 10 minutes. Urea/DTT 10 M solution was added and homogenized. The samples were incubated at 95°C for 3 minutes, sonicated and stored at −80°C. The total protein concentration was quantified using the bicinchoninic acidic (BCA) protein assay (PierceTM BCA Protein Assay Kit – Thermo Scientific) or the Bradford assay (NZYTech) according with the manufacture’s instruction. Samples were denaturized by adding Laemmli SDS sample buffer (4X) (Alfa Aesar by Thermo Fisher Scientific) and heated at 95°C for 5 min. RIPA buffer was used to minimize variations in protein extracts. Samples were stored at −20°C until western blot analysis.

### Quantitative reverse-transcription PCR

RNA was isolated using the NZYTech™ RNA Isolation kit (NZYtech) according to the manufacturer’s instructions. RNA was reverse transcribed with the iScriptTM cDNA Synthesis kit (Bio-Rad) to generate complementary DNA. Quantitative reverse-transcription PCR (q-RT-PCR) was performed on a CFX96 Touch Real-Time PCR Detection System (Bio-Rad) using the SsoAdvancedTM SYBR® Green SuperMix kit (Bio-Rad) and Invitrogen (ThermoFisher Scientific) designed-primers for the Human *CYP46A1* (Fwd: GCAGCGGAGTCATAGACC; Rev: CAGCAGCATACTGGTCTCCA), Mouse *Cyp46a1* (Fwd: TCCTCTCCTGTTCAGCACC; Rev: CAGCTTGGCCATGACAACT) and *Npy*^16^. Expression levels of target genes were normalized to the expression of the housekeeping gene Mouse *HPRT* (Fwd: GCTTACCTCACTGCTTTCCG; Rev: CATCATCGCTAATCACGACGC). Controls were used to exclude the possibility of DNA or RNA contamination.

### Western Blotting

Protein samples were loaded in a 12% polyacrylamide gel. After SDS-PAGE for 80 V for 30 minutes and 120 V for 1 hour. Later gels were blotted onto a methanol 99,9% (Fisher Chemical) activated Polyvinylidene fluoride (PVDF) (Merck Millipore) membrane at a constant current of 500 mA for 4 hours at 4°C. Later membranes were blocked in 5% bovine serum albumin (BSA) in TBS-T for 1 hour and probed with the following antibodies: PPAR-γ (1:1000, Cell Signaling Technology, #81B8) and β-Actin (1:5000, Sigma-Aldrich, A5316). Antibodies were diluted in 5% BSA in TBS-T. Next, the membranes were incubated with horseradish peroxidase-conjugated secondary antibodies and later analyzed using a ChemiDoc^TM^ Imaging System (Bio-Rad, California, USA).

### Tissue processing and Hematoxylin-Eosin staining

Organs and tissues were fixed in a 4% Formaldehyde solution, stabilized with methanol fixative solution and positioned in tissue processing cassettes (Labor Spirit). Dehydration of tissues was achieved through 70% Ethanol (Fisher Chemical) for 1 hour, two series of 95% Ethanol for 45 minutes each and two series of 100% Ethanol for 1 hour each. After, the clearing was performed with two series of xylene incubations (Fisher Chemical) for 1 hour each and wax infiltration with two series of paraffin (Luso Palex) each at 56°C for 1 hour. The cassettes were set in embedding molds (Tebu-bio), to form a block, at a chosen orientation and filled with liquid paraffin at 56°C then let to cool down and solidify. The block was removed from the mold and stored at room temperature until use.

The paraffin blocks were cut into paraffin sections on a HM 325 Rotary Microtome (Thermo Fisher Scientific) at room temperature. The paraffin blocks that contained adipose tissue were cut into 4-5 μm paraffin sections and the liver and pancreas were cut into 3-4 μm sections then placed into microscopy slides and stored at room temperature. Hematoxylin-Eosin staining was performed according to the manufacturer guidelines (Merck Millipore). The microscope slides were deparaffinized by immersing them in two series of xylene for 3 minutes and 2 minutes, respectively. Then, they were rehydrated through a series of solutions, starting with 100% Ethanol for 4 minutes, followed by 95% Ethanol for 2 minutes. Finally, the slides were rinsed in two series of distilled water for 30 seconds each. The sections were stained with a modified Hematoxylin solution modified (Merck Millipore) for 30 seconds followed by 2 washes in distilled water for 2 and 1 minutes, respectively. They were then counterstained with a 0.5% aqueous eosin Y solution (Acros Organics) for 1 minute; followed by two washes in distilled water for 1 minute each, and two series of xylene for 2 minutes. After drying, the sections were mounted using Richard-Allan Scientific Mounting Medium (HM325, ThermoFisher Scientific) and covered with microscopy slide coverslips.

### Microscopy

Images were acquired in a Zeiss Axio Imager Z2 microscopy (Carl Zeiss) at ×10 or ×20 magnification using an AXIOCAM-ICC3 camera associated with the AxioVision software (Carl Zeiss). The quantification of the images obtained were performed with Fiji. The images were analyzed, in a blinded fashion, to calculate the area of the adipocytes, and respective mean area (both made with tools from Fiji) was determined.

Pancreas images were analysed in pancreatic islets area and liver samples were analyzed to obtain a histological score, relative to the presence of steatosis or other hepatic anomalies. Ballooning was scored from 0-2: 0 for physiological hepatocytes (cuboidal shape), 1 for presence of clusters of hepatocytes with a rounded shape and at last, 2 for the same grade as 1, but with enlarged hepatocytes (at least 2-fold) according to previous descriptions^17^.

### Glucose and Insulin tolerance tests (GTT and ITT)

Mice from subject to a glucose tolerance test (GTT). Briefly, animals were subjected to overnight starvation. After this period, blood was collected to measure the fasting blood glucose levels (timepoint 0). Blood glucose levels measurements were performed using the FreeStyle Precision Neo glucometer and FreeStyle Precision strips (Abbot). After the measurement of the fasting blood glucose level, mice were injected intraperitoneally with a 20% glucose solution (Fisher Chemical - G/0500/60), dissolved in saline solution, 0,9 % NaCl. The injection volume was calculated according to the formula^19^: *Injection volume = Body weight (g) 10 μL 20% glucose solution*. Blood glucose levels were measured at 5, 10, 15, 30, 60, 90 and 120 min after the intraperitoneal injection of glucose solution. The measured levels were used to perform the calculation of the area under the curve (AUC). For this, the blood glucose measurements were converted into natural logarithm (Ln) and the total peak area was calculated using xy analysis. Two values were calculated: the total AUC, from 0 until 120 min, and the AUC between 5 and 60 min.

The Insulin Tolerance Test (ITT) was also conducted after a starvation period (4 hours). Blood was collected to measure the fasting blood glucose levels (timepoint 0). The blood glucose levels measurements were performed using the FreeStyle Precision Neo glucometer and FreeStyle Precision strips (Abbot). After the measurement of the fasting blood glucose level, mice were injected intraperitoneally with a 4 mg/mL human insulin solution (GibcoTM) in a phosphate-buffered saline solution. The injection volume was calculated using the following formula: *Injection volume = Body weight (g) 7.5 μL of 4 mg/mL insulin solution*. The blood glucose levels were measured at 5, 10, 15, 30, 60, 90 and 120 min after the insulin intraperitoneal injection. At the end of the ITT test, the animals were injected with a 20% glucose solution to revert insulin effect and to prevent animal death. The blood glucose measurements were then converted into natural logarithm (Ln) and the slope was calculated using linear regression and multiplied by 100 to obtain the constant for glucose clearance (kITT), per minute (%/min) obtained during the insulin tolerance test ^20^.

### DNA extraction from Fecal samples and bacterial taxa quantification by qPCR

For DNA extraction 100-240 mg of feces of mice were used. The DNA extraction was performed using the QIamp PowerFecal DNA Kit (Qiagen) according to the manufacturer’s instructions with a slight modification on the lysis step to reduce the degradation of the extracted DNA. For this, instead of using a beadbeater for fecal homogenization and bacterial lysis, temperature treatment was employed as an alternative. This was done after the addition of the C1 reagent. Samples were homogenized by vortex for 5 seconds followed by water bath incubation at 70°C for 5 min, following a new vortex step for 5 seconds and a final water bath exposure at 70°C for 5 min. The extracted DNA was eluted in C6 solution and maintained at −20°C. Each sample of DNA was quantified using a Nanodrop (Fisher Scientific) and its integrity was assessed by agarose gel electrophoresis. For qPCR of each bacterial taxa the 16S rRNA gene was amplified with specific primers from the fecal samples. Each amplified fragment was inserted into the pCR ™ 2.1-TOPO cloning vector (Invitrogen) and transformed in *Escherichia coli* MACH1 competent cells. After colony selection plasmids were extracted and purified for identification of each bacterial group of interest using the specific forward and reverse primers. After the sequencing validation the plasmids were used on quantitative real-time PCR.

### Quantitative Polymerase chain Reactions (qPCR)

Standard curves were constructed for each bacterial group by using dilutions of plasmid DNA. The mass of each amplified fragment was calculated using the following formula: *Mass of each fragment (ng)* = size of the fragment (bp)×(1.096×10^−21^). The mass of each fragment was used to calculate the copy numbers per μL in each dilution using the following formula, and then converted to logarithmic scale:

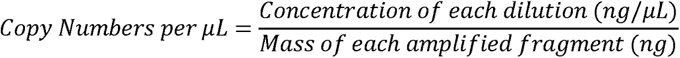

To perform the bacterial quantifications, SsoFastTM kit Eva Green Supermix (BioRad) was used and the PCR reaction. The copy numbers of each group present in each sample were inferred from each constructed standard curve. The amplification efficiency of each reaction was calculated using the formula below:

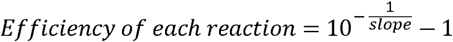

For analysis of fecal samples the following previously validated primers were used: *Bacteroidetes*^18^ (Fwd:CATGTGGTTTAATTCGATGAT; Rev: AGCTGACGACAACCATGCAG), *Firmicutes*^19^ (Fwd: GGAGYATGTGGTTTAATTCGAAGCA; Rev: AGCTGACGACAACCATGCAC), *Actinobacteria*^19^ (Fwd: TGTAGCGGTGGAATGCGC; Rev: AATTAAGCCACATGCTCCGCT), *Tenericutes*^19^ (Fwd: ATGTGTAGCGGTAAAATGCGTAA; Rev: CMTACTTGCGTACGTACTACT), *Betaproteobacteria*^19^ (Fwd: AACGCGAAAAACCTTACCTACC; Rev: TGCCCTTTCGTAGCAACTAGTG;) *Delta and Gammaproteobacteria*^19^ (Fwd: GCTAACGCATTAAGTRYCCCG; Rev: GCCATGCRGCACCTGTCT), *16S rRNA* gene^19^ (Fwd: AGAGTTTGATCCTGGCTCAG; Rev: AAGGAGGTGWTCCARCC), Universal^19^ (Fwd: AAACTCAAAKGAATTGACGG; Rev: CTCACRRCACGAGCTGAC) and *dsrB* gene^20^ (CAACATCGTYCAYACCCAGGG; GTGTAGCAGTTACCGCA).

### Stereotaxic surgery

rAAVs were injected into the arcuate nucleus of the mouse hypothalamus at the following coordinates relative to Bregma: − 1.65 mm anteroposterior, ± 0.5 mm medio-lateral, −5.8 mm dorsoventral. A total of 1×10^9^ v.g. (viral genomes) in a total volume of 1.5 µL was injected per hemisphere at 300 nL/min using a 10 mL-Hamilton syringe attached to a Quintessential Stereotaxic Injector (QSI™) (Stoelting Company) as previously performed^21^. Following each injection, the needle was left in place for 5 minutes to ensure proper diffusion. During behavioral testing, the experimenter was blinded to the identity of the virus injected into each mouse. Behavioral experiments started 4 weeks after rAAVs delivery.

### Statistical analysis

Each set of experiments contained mice injected with control or experimental viruses and were randomized per cage (i.e., each cage of four mice contained mice injected with control or experimental viruses). After stereotaxic surgery and until the end of each experiment, the authors were blind to the experimental group of each mouse. For normally distributed data sets, two-tailed unpaired Student’s t tests were used to compare the two groups. In the case of datasets which compare more than once a one- or two-way ANOVA was used followed by appropriate multiple comparison post hoc tests to control, for multiple comparisons as specified. In the case of a non-Gaussian distribution, two-tailed Mann-Whitney tests were used to compare two distinct groups, or a Kruskal-Wallis test followed by Dunńs post hoc test to compare more than 2 groups. Principal component analysis (PCA) was performed using R (packages vegan version 2.6-6.1 and ggplot2 version 3.5.1) in all bacterial groups. Nested analysis was performed on single microbiota data using a nested Student’s t test. The sample size was determined based on similar experiments carried out in the past and in literature. Statistical analysis was performed using GraphPad Prism, version 11. For behavioral experiments the investigators were blind to group allocation during data collection and analysis. The exact sample size is depicted by individual dots in figures.

## Results

### Diet-induced obesity promotes hypothalamic CYP46A1 decrease

To investigate the potential role of hypothalamic cholesterol metabolism in diet-induced obesity, C57BL/6J mice were assigned to two distinct dietary regimens: a standard chow diet and a high-fat diet (HFD) (Figure 1A). Mice were sacrificed and the hypothalamus was microdissected to analyze mRNA levels of the central nervous system-specific Cyp46a1 enzyme. We found that Cyp46a1 mRNA levels were reduced in mice that underwent a HFD diet compared to those that underwent chow diet (Figure 1B). This observation suggests that hypothalamic CYP46A1 levels may contribute to the metabolic adaptations associated with high-calorie intake, potentially leading to obesity development. To test this hypothesis, we generated recombinant adeno associated viruses (rAAV) harboring expression cassettes to knockdown (shCyp46a1) or overexpress CYP46A1 (CYP46A1) in the arcuate nucleus (ARC) of the hypothalamus (Figure 1A-B, Supplementary Figure 1A-B). These rAAVs were delivered to the ARC of mice that underwent chow and HFD diets, and food and water consumption, as well as body weight, were monitored for 6 weeks post-surgery. Control mice that underwent HFD displayed increase food intake and total body weight gain. Mice injected with shCyp46a1 on the chow diet showed significant increases in food and water consumption overtime compared to controls (Figure 1C-F). Conversely, HFD shCyp46a1 injected mice showed no changes in these parameters. However, both groups exhibited a significant increase in body weight in conditions of shCyp46a1 (Figure 1G-I). These findings indicate that reduced CYP46A1 levels in the hypothalamus fosters increased feeding behaviors and contribute to an obesity phenotype.

**Figure 1.**
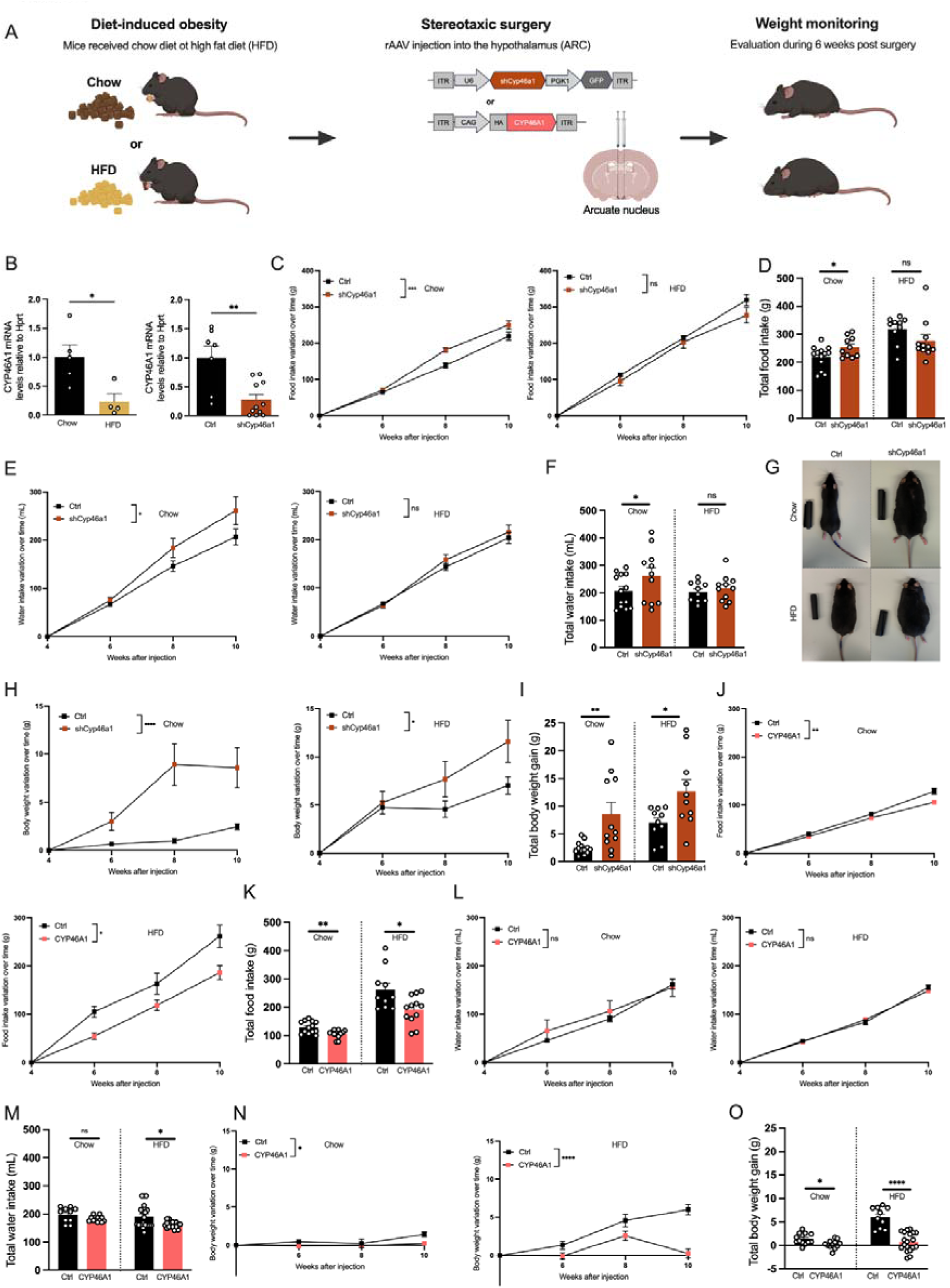
High-fat diet-associated decrease in CYP46A1 levels leads to obesity. (**A**) Experimental scheme. (**B**) Mice underwent chow or high fat diet (HFD) for 4 weeks their hypothalamus was dissected and mRNA isolated to perform RT-qPCR for CYP46A1 mRNA levels (N=4-5; N=7-11). Mice were stereotaxically injected in the arcuate nucleus with rAAVs encoding an shRNA CYP46A1. 4-weeks post-surgery mice were evaluated every two weeks for food (**C-D**) or water intake (**E-F**) variation and (**G-I**) body weight (N=10-13). Mice were stereotaxically injected in the arcuate nucleus with rAAVs encoding for CYP46A1. 4-weeks post-surgery mice were evaluated every two weeks for food (J-K) or water intake (L-M) variation and (N-O) body weight (N=12-13). Dots represent individual mice. Data represents mean ± standard error of the mean (SEM). For time-course experiments a two-way repeated measure ANOVA was used. One-way ANOVA; Dunnett’s, Šídák’s or Fisher’s test; Ns, nonsignificant, *p<0.05, ** p<0.01, **** p<0.0001.

Next, we evaluated whether increasing CYP46A1 levels would yield opposite results. Both chow and HFD CYP46A1-overexpressing mice showed decreased food and water intake (Figure 1J-M). As expected, body weight was consistently reduced under CYP46A1 overexpression conditions (Figure 1N-O). These changes were not due to virus-specific toxicity affecting hypothalamic function (Supplementary Figure 1C-D). Altogether these results show that modulation of CYP46A1 levels in the ARC alters diet-induced obesity phenotypes.

### CYP46A1 levels modulate adipocyte hypertrophy and PPAR-**γ** levels

During diet induced obesity there is adipocyte size increase due to the accumulation of lipids in the form of triglycerides ^22,23^. The changes in body weight observed upon modulating CYP46A1 levels prompted us to investigate whether these were associated with alterations in adipocyte size. Mice on both diets were stereotaxically injected with rAAVs to either downregulate or overexpress CYP46A1 expression. Mice were then sacrificed, and white adipose tissue was collected, weighed and processed for histological and biochemical analysis (Figure 2A). As expected, HFD lead to adipocyte hypertrophy in control conditions. We found that in both dietary regimens, mice with reduced CYP46A1 levels exhibited a higher frequency of adipocytes with larger areas, along with an overall increase in WAT weight (Figure 2B-D). Next, we performed a similar analysis to animals overexpressing CYP46A1 levels in the ARC. In contrast, mice overexpressing CYP46A1 in the ARC showed a higher frequency of adipocytes with smaller areas and a general reduction in WAT weight (Figure 2E-G). These results were not due to virus-specific toxicity (Supplementary Figure 2A-B). The observed alterations in WAT morphology are highly suggestive of a shift in cellular metabolism. To test this hypothesis, we analyzed the protein levels of the peroxisome proliferator-activated receptor gamma (PPAR-γ), a known regulator of adipogenesis and a potent modulator of whole-body lipid metabolism and of insulin sensitivity^24^. shCyp46a1 injected mice displayed reduced PPAR-γ levels specifically under an HFD, compared to controls (Figure 2H-I). In contrast, mice overexpressing CYP46A1 showed an overall increase in PPAR-γ levels (Figure 2H-I). Altogether, this set of results indicate that CYP46A1 levels modulate diet-induced obesity and adipocyte hypertrophy.

**Figure 2.**
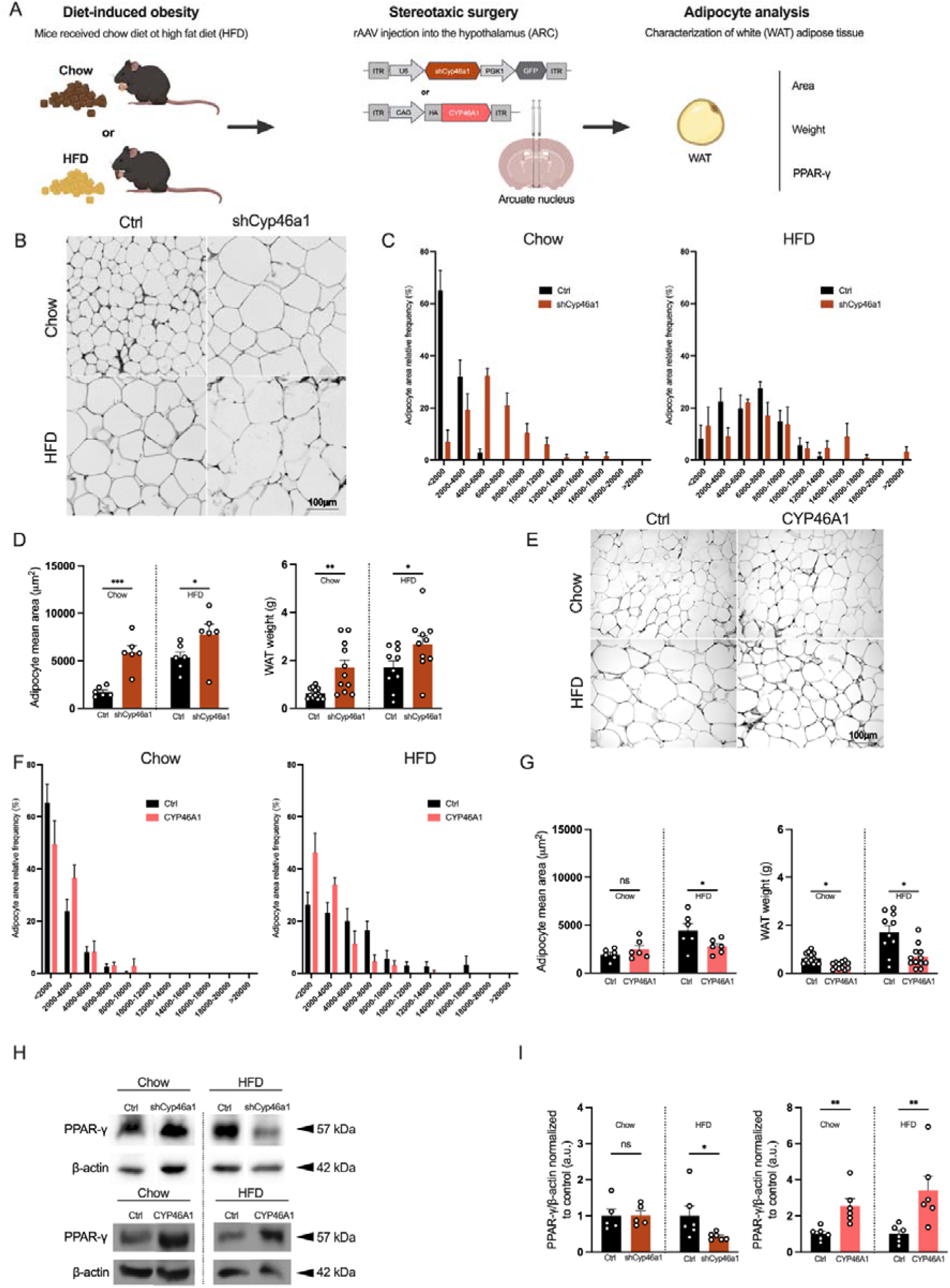
Hypothalamic CYP46A1 levels mediate adipocyte hypertrophy. (**A**) Experimental scheme. (**B**) Mice underwent chow or high fat diet (HFD) for 4 weeks and were stereotaxically injected with AAVs expressing a CYP46A1-shRNA (**A-D**) or CYP46A1 (**E-G**) in the arcuate nucleus of the hypothalamus. White adipose tissue (WAT) was isolated for histological analysis of adipocyte area (N=6) and weight (N=10-13). (**H-I**) Protein was isolated for western blot analysis from HFD for detection of the peroxisome proliferator-activated receptor gamma (PPAR-γ) (N=5-6). Dots represent individual mice. Data are shown as mean ± standard error of the mean (SEM). One-way ANOVA; Dunnett’s, Šídák’s or Fisher’s test; Ns, nonsignificant, *p<0.05, ** p<0.01, *** p<0.001. Scale bar =100 μm.

### CYP46A1 modulation promotes organism-wide alterations associated with diet-induced obesity

Diet-induced obesity is known to disrupt whole-body metabolic function, affecting multiple organs. This includes insulin resistance, which reduces hypothalamic expression of orexigenic neuropeptides like neuropeptide Y (NPY) that regulate appetite, as well as liver inflammation and increased pancreatic islet size ^25,26^. We hypothesized that these features could be mediated by CYP46A1. To evaluate this possibility, we dissected and collected the hypothalamus, liver, spleen and pancreas from mice expressing shCyp46a1, CYP46A1 or controls, that underwent chow or HFD regimens (Figure 3A). As expected, we found that HFD reduced the levels of NYP in the ARC compared to control chow mice ^27^(Figure 3B). Interestingly, we found that this effect was fully recapitulated under shCyp46a1 conditions and reversed in CYP46A1-overexpressing mice (Figure 3B). Additionally, reduction of ARC CYP46A1 levels led to significant alterations in the total weight of the liver and spleen (Figure 3C-D, Supplementary Figure 3A-B). We then conducted a histological analysis of hepatocyte morphology (Figure 3E-F). As expected, control animals that underwent HFD showed hepatocyte enlargement (Figure 3E-F). We found that shCyp46a1-injected mice undergoing chow diet displayed enlarged hepatocytes, like the controls under HFD (Figure 3E-F). Conversely, overexpressing CYP46A1 prevented diet-induced hepatocytes enlargement. We then assessed hepatic steatosis by quantifying the area of lipid droplets in liver tissue. These experiments showed that reducing CYP46A1 levels promoted hepatic steatosis in mice on a chow diet, similar to the effects observed in control mice that underwent HFD (Figure 3G). Hepatic steatosis was prevented in CYP46A1-overexpressing mice that underwent HFD.

**Figure 3.**
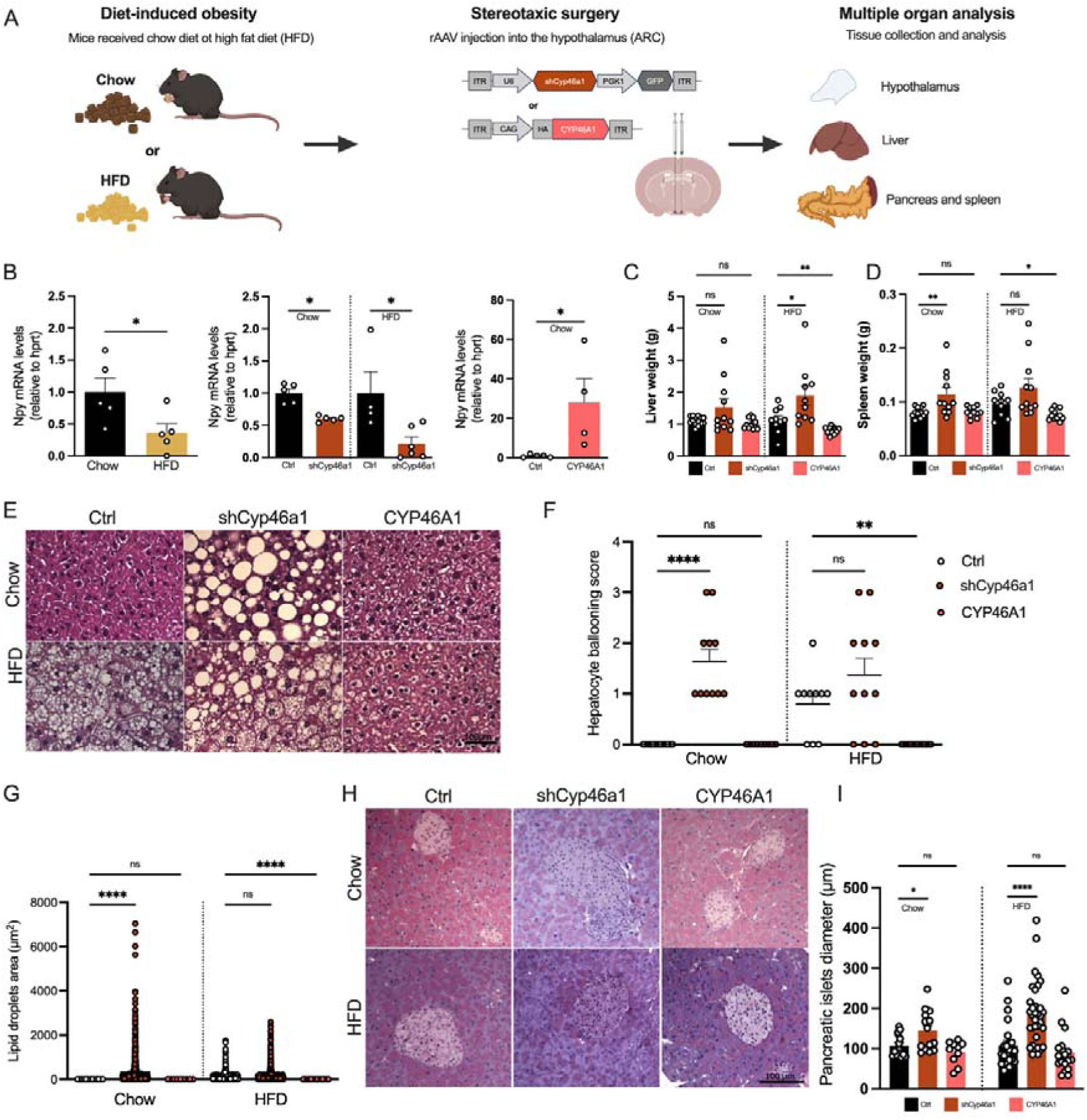
Hypothalamic, hepatic and pancreatic adaptations to diet-induced obesity are mediated by CYP46A1 levels. (**A**) Experimental scheme. (**B**) Mice underwent chow or high fat diet (HFD) for 4 weeks and were stereotaxically injected with AAVs expressing a CYP46A1-shRNA or CYP46A1 in the arcuate nucleus of the hypothalamus. Hypothalami were dissected and mRNA isolated to perform RT-qPCR for NPY mRNA levels (N=4-6). Organs were collected for weight evaluation as raw measure of fat accumulation in tissues, particularly (**C**) liver (N=10-13) and (**D**) spleen (N=10-12). (**E**) Histological analysis was performed on liver tissue to evaluate alterations in (**F**) hepatocyte morphology (ballooning) (N=9-14) as well as (**G**) the size of lipid droplets (N=9-14). (**H-I**) Pancreatic tissue was analyzed to evaluate the total diameter of islets of Langerhans which are associated with insulin-secretion (N=10-34). Dots represent individual mice, except in (**G**) which represent individual droplets. Data are shown as mean ± standard error of the mean (SEM). Unpaired two-tailed Student’s T test; One-way ANOVA; Dunnett’s, Šídák’s or Fisher’s test, Ns, nonsignificant, *p<0.05, ** p<0.01, **** p<0.0001. Scale bar =100 μm.

Diet-induced obesity can lead to insulin resistance, triggering compensatory adaptations in the pancreas, such as an increase in the size of pancreatic islets^28^. To investigate whether CYP46A1 expression contributes to changes in pancreatic histology, we examined pancreatic islet morphology. Control animals that underwent HFD had increased pancreatic islet size (Figure 3H-I). We found that reducing the levels of CYP46A1 mimicked the diet-induced enlargement of pancreatic islets (Figure 3H-I) in chow conditions. These results indicate that modulation of CYP46A1 levels in the ARC plays a key role in the hypothalamic, hepatic, and pancreatic pathological features associated with diet-induced obesity.

### Diet-associated hyperglycemia and insulin resistance are dependent on hypothalamic Cyp46a1 levels

Our observation that hypothalamic Cyp46a1 mediates obesity-related changes in multiple organs, including the pancreas, suggests it may play a role in regulating glucose metabolism. To investigate this hypothesis, we measured blood glucose levels in mice under baseline conditions and after intraperitoneal administration of a 20% glucose solution (Figure 5A). The glucose tolerance test (GTT) revealed that control animals that underwent HFD were hyperglycemic compared to chow fed mice (Figure 5B-C). These results suggest that these mice may have insulin resistance, leading to reduced glucose uptake over time. To confirm this possibility, mice were subjected to an insulin tolerance test (ITT), where blood glucose levels were measured under baseline conditions and after intraperitoneal injection with a 4 mg/mL human insulin solution (Figure 5A). This experiment confirmed reduced glucose uptake in response to insulin in control mice that underwent HFD, compared to chow-fed mice (Figure 5D-E). Next, we performed similar experiments on shCyp46a1 and CYP46A1-overexpressing animals. We found that reducing CYP46A1 levels increased glucose levels in chow, but not in those on an HFD condition compared to controls (Figure 5F-E). Consistent with this, we found that reducing CYP46A1 levels resulted in a reduced plasma glucose disappearance rate, a clear indication of insulin resistance (Figure 5I-J). Based on these findings, we tested whether hyperglycemia and insulin resistance could be alleviated by CYP46A1 overexpression. The GTT revealed that CYP46A1-injected mice showed a selective reduction in glucose levels in HFD conditions (Figure 5L-N). Moreover, we also detected reduced glucose levels in these animals during the ITT, although the plasma glucose disappearance rate remained similar between groups (Figure O-Q). Altogether, these results show that hypothalamic CYP46A1 levels modulate circulating glucose metabolism and uptake.

**Figure 4.**
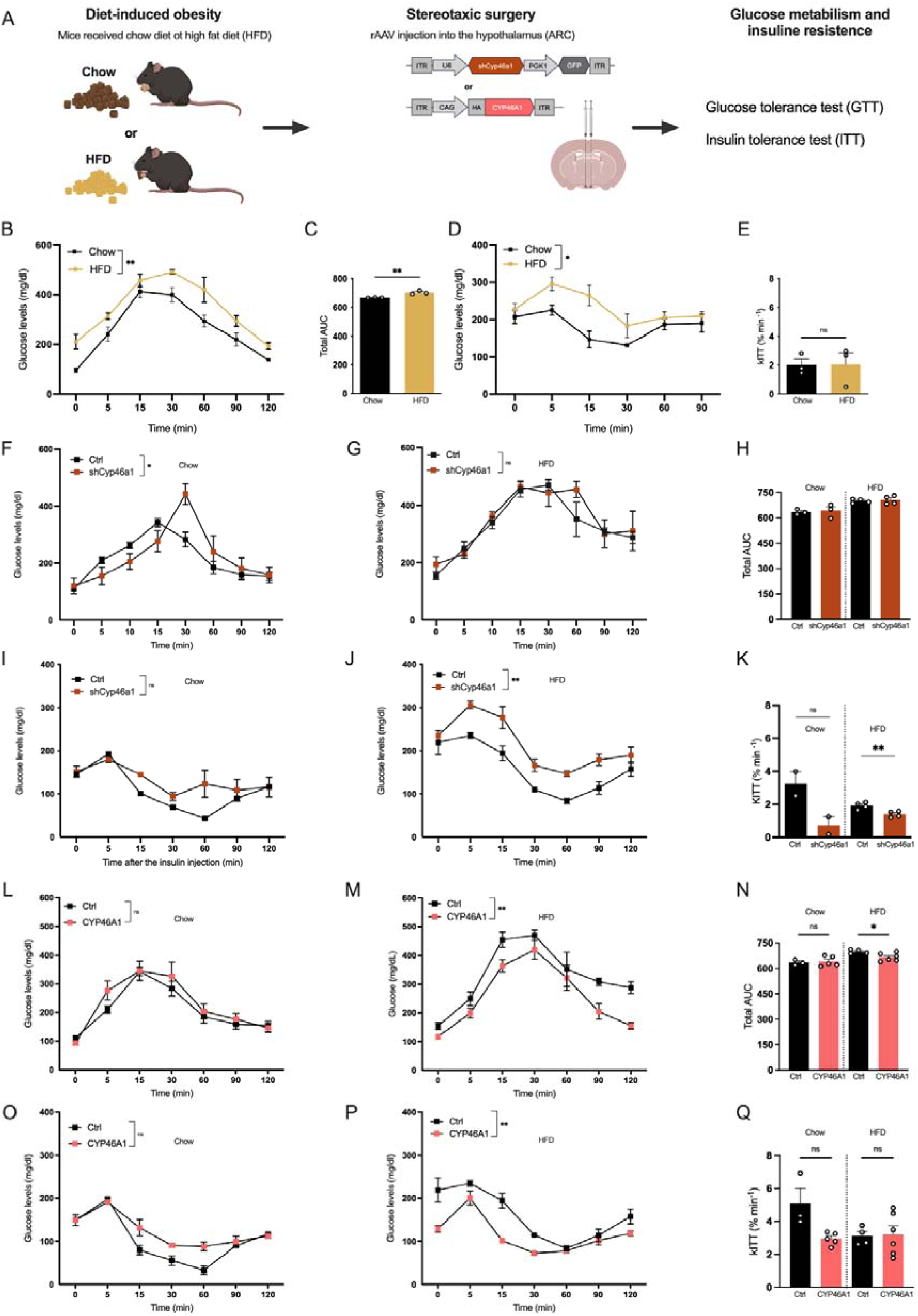
Hypothalamic Cyp46a1 levels regulate hyperglycemia and insulin insensitivity. (**A**) Experimental scheme. Mice underwent chow or high fat diet (HFD) for 4 weeks and were stereotaxically injected with AAVs expressing a CYP46A1-shRNA or CYP46A1 in the arcuate nucleus of the hypothalamus. Mice were subjected to both diets underwent a (**B-C**) glucose tolerance test (GTT) or (**D-E**) insulin tolerance test (ITT) (N=3) to evaluate the area under the curve (AUC) or plasma glucose disappearance rate (kITT). (**F-K**) CYP46A1-shRNA (N=2-4) or (**L-Q**) CYP46A1 (N=3-6) injected animals were also evaluated in both tests. For time-course experiments a two-way repeated measure ANOVA was used. One-way ANOVA; Dunnett’s, Šídák’s or Fisher’s test; Ns, nonsignificant, *p<0.05, ** p<0.01.

**Figure 5.**
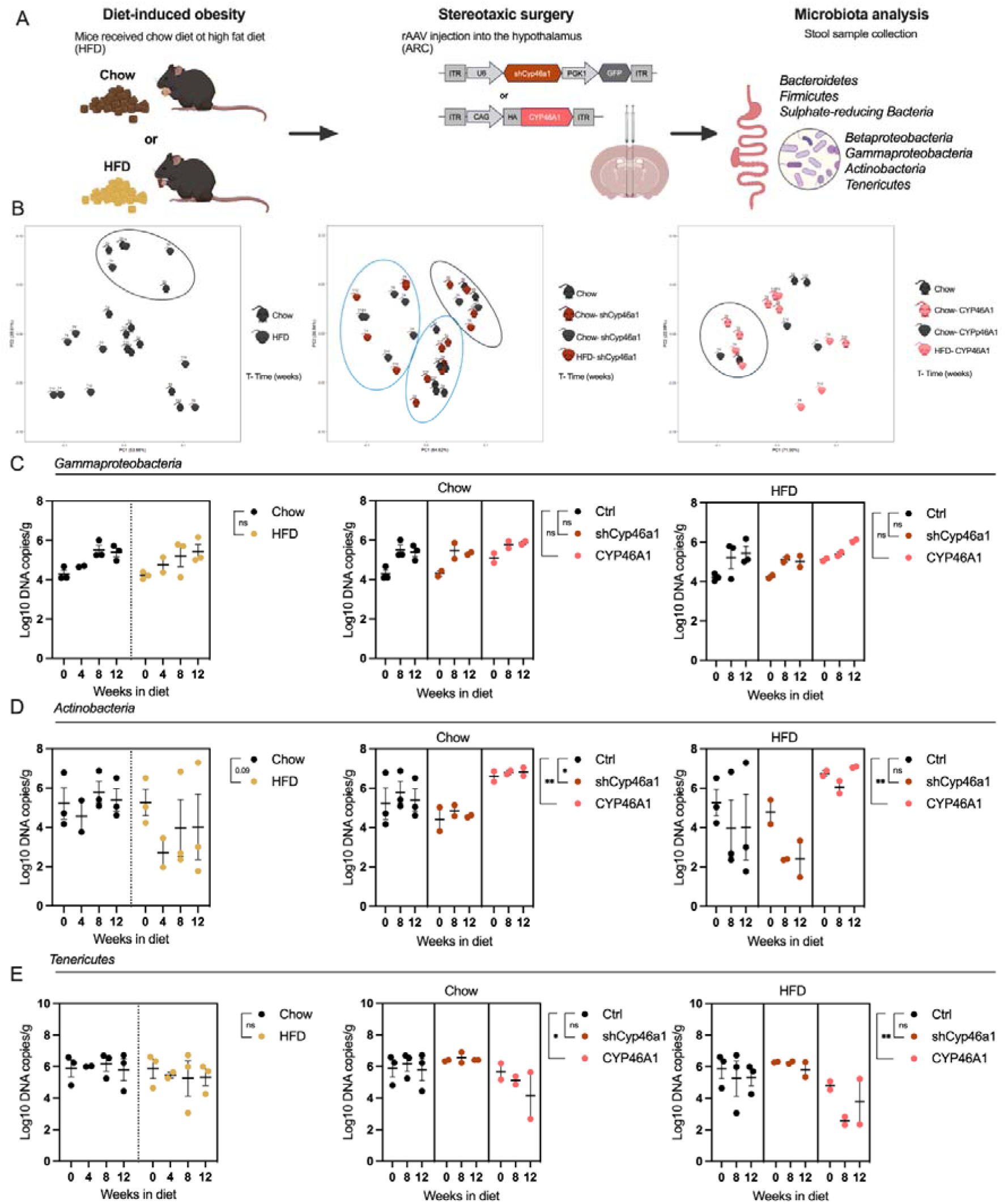
Hypothalamic Cyp46a1 levels mediate diet-associated microbiota composition (**A**) Experimental scheme. Mice underwent chow or high fat diet (HFD) for 4 weeks and were stereotaxically injected with AAVs expressing a CYP46A1-shRNA or CYP46A1 in the arcuate nucleus of the hypothalamus. Stool samples from mice were collected and the counts of bacterial taxa by qPCR as a measure of gut microbiota: *Bacteroidetes, Firmicutes, Sulphate-reducing Bacteria, Betaproteobacteria, Gammaproteobacteria, Actinobacteria and Tenericutes*. (**B**) Principal component analysis of the seven Phyla according to diet or CYP46A1 modulation was performed. Evaluation of by qPCR according to diet (Chow vs HFD, left) (N=11) or CYP46A1 modulation (shCyp46a1 or CYP46A1 overexpression) (N=6-9) of Phyla levels: **(C)** *Gammaproteobacteria*, **(D)** *Actinobacteria* and **(E)** *Tenericutes*. Nested Student’s t test, Ns, nonsignificant, *p<0.05, ** p<0.01.

### Hypothalamic Cyp46a1 levels mediate gut diet-associated microbiota changes

It is now well established that obesity is a multifactorial disorder influenced not only by the balance between energy intake and expenditure, but also by other causative factors such as the gut microbiome^29,30^. In the next set of experiments, we quantified the abundance of several well-characterized intestinal bacterial groups, in particular *Bacteroidetes*, *Firmicutes*, Sulphate-Reducing Bacteria (SRB), *Betaproteobacteria*, Delta-and *Gammaproteobacteria*, *Actinobacteria* and *Tenericutes.* We collected stool samples from mice that underwent both dietary regimens and compared the abundance of these bacteria over time (Figure 5A). Principal component analysis (PCA) revealed a single cluster at timepoint 0, indicating that bacterial load and composition were similar across all groups at the onset of the diets (Figure 5B). Moreover, we observed clustering defining Chow and HFD groups, specifically for mice treated with shCyp46a1. Next, we individually evaluated each of the seven bacterial taxa. This analysis showed that neither diet nor Cyp46a1 manipulation affected the abundance of *Bacteroidetes*, *Firmicutes*, *SRB*, *Betaproteobacteria* and *Delta*-*and Gammaproteobacteria* (Figure S4 and Figure 5C).

We observed a trend toward decreased *Actinobacteria* levels upon HFD treatment compared to Chow mice (Figure 5D). This decrease was recapitulated in shCyp46a1 mice while the opposite effect was observed in CYP46A1 mice undergoing Chow diet (Figure 5D). These results suggest that the predominance of *Actinobacteria* in the gut of mice undergoing HFD is modulated by hypothalamic *Cyp46a1* levels. No diet-specific changes were observed in *Tenericutes* levels, although CYP46A1-overexpressing mice exhibited a reduction in this bacterial group across both diets (Figure 5E).

### Hypothalamic Cyp46a1 levels regulate anxiety-like behavior, motor activity and cognitive function associated with diet-induced obesity

The broad metabolic alterations resulting from hypothalamic modulation of Cyp46a1 levels led us to hypothesize that these changes might also affect behavior. To this end, we conducted open field and Y-maze tests since diet-induced obesity has been associated with anxiogenic-like behavior and cognitive dysfunction in mice^31^. Mice that underwent both dietary regimens were stereotaxically injected with rAAVs encoding CYP46A1-shRNA or CYP46A1 and subsequently underwent both behavioral tests (Figure 6A). The open field test was conducted at 4- and 8-weeks post-injection during the light phase, and 8-weeks post-surgery in the dark phases, as obesity is known to impact mouse performance differently during day and night ^32,33^. During the inactive phase (day), mice with CYP46A1 knock-down spent less time in the center of the open field and showed increased grooming behavior, with no changes in locomotion compared to controls (Figure 6B-E). Similarly, CYP46A1-overexpressing mice showed increased grooming behavior, but no alterations were found in time spent in the center of the arena, nor in locomotion activity (Figure 6F-I). Next, we evaluated the same parameters during the active phase (night) and found that CYP46A1-shRNA injected mice spent more time in the center of the arena, while CYP46A1 injected mice displayed reduced exploration of this region (Figure 6J). Moreover, CYP46A1 knock-down and overexpression led to increased or decreased immobility, respectively (Figure 6K). Consistent with the active phase results, animals with both manipulations exhibited increased grooming behavior (Figure 6L). Animals with reduced CYP46A1 levels displayed decreased locomotion compared to controls (Figure 6M). Altogether, these results revealed that the obesity-associated reduction of hypothalamic CYP46A1 levels decreases locomotor activity and alters anxiety-like behavior. Next, we evaluated working memory performance in the Y-maze paradigm. Over time, animals with reduced levels of CYP46A1 showed fewer spontaneous alternations, indicating impaired working memory (Figure 6N). In contrast, mice overexpressing CYP46A1 showed an increase in spontaneous alternations, suggesting improved cognitive abilities (Figure 6N). Together, these findings demonstrate that hypothalamic CYP46A1 levels bidirectionally modulate anxiety-related behaviors and working memory performance in the context of diet-induced obesity.

**Figure 6.**
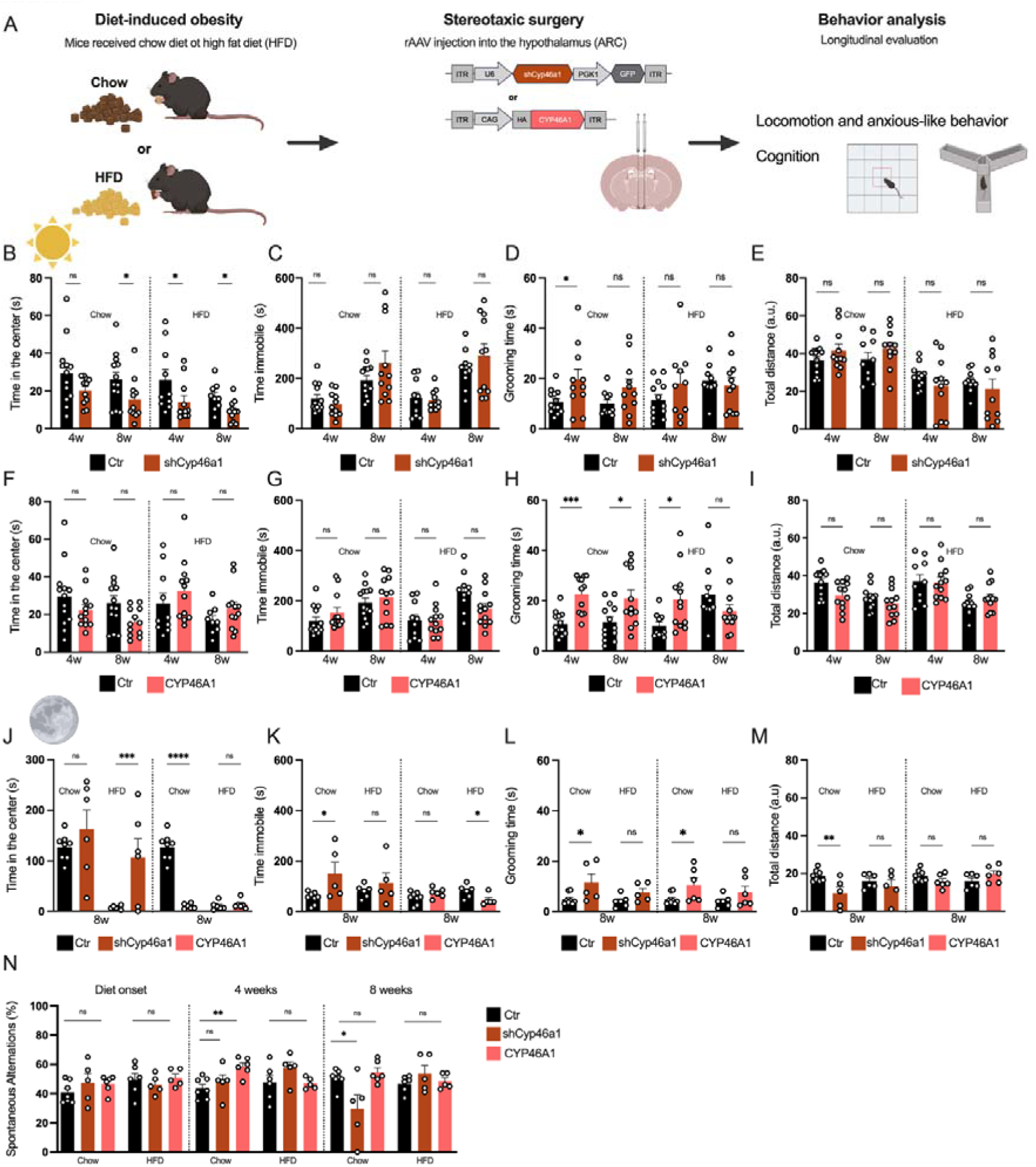
Hypothalamic Cyp46a1 levels impact anxious-like behavior, locomotion and cognition (**A**) Experimental scheme. Mice underwent chow or high fat diet (HFD) for 4 weeks and were stereotaxically injected with AAVs expressing a CYP46A1-shRNA or CYP46A1 in the arcuate nucleus of the hypothalamus. Mice were subjected to a battery of behavior paradigms 4 and 8 weeks post-surgery, the open field test during daytime (**B-I**) (N=10-13) and nighttime (**J-M**) (N=6-8) periods to evaluate anxiety-like behavior and locomotion and the Y-maze test for working memory analysis (**N**) (N=5-7). One-way ANOVA; Dunnett’s, Šídák’s or Fisher’s test; Ns, nonsignificant, *p<0.05, ** p<0.01, *** p<0.001 and **** p<0.0001.

## Discussion

In this study, we uncovered a novel mechanism regulated by the hypothalamus during high-caloric-associated obesity. In particular, the CYP46A1, a key enzyme in cholesterol metabolism, emerged as a potential link between high-caloric intake and hypothalamic control of whole-body metabolism. We demonstrated that high-caloric intake reduces hypothalamic CYP46A1 levels. Using genetic approaches to either model the decrease or overexpression of CYP46A1, we showed that CYP46A1 levels bidirectionally regulate obesity-related phenotypes, including adipocyte size, glucose homeostasis and insulin sensitivity. Furthermore, our findings revealed that this enzyme also influences gut microbiota composition, as well as cognitive and motor behaviors, highlighting its multifaceted role in metabolic and neurological regulation (for graphical summary please see Figure 7).

**Figure 7.**
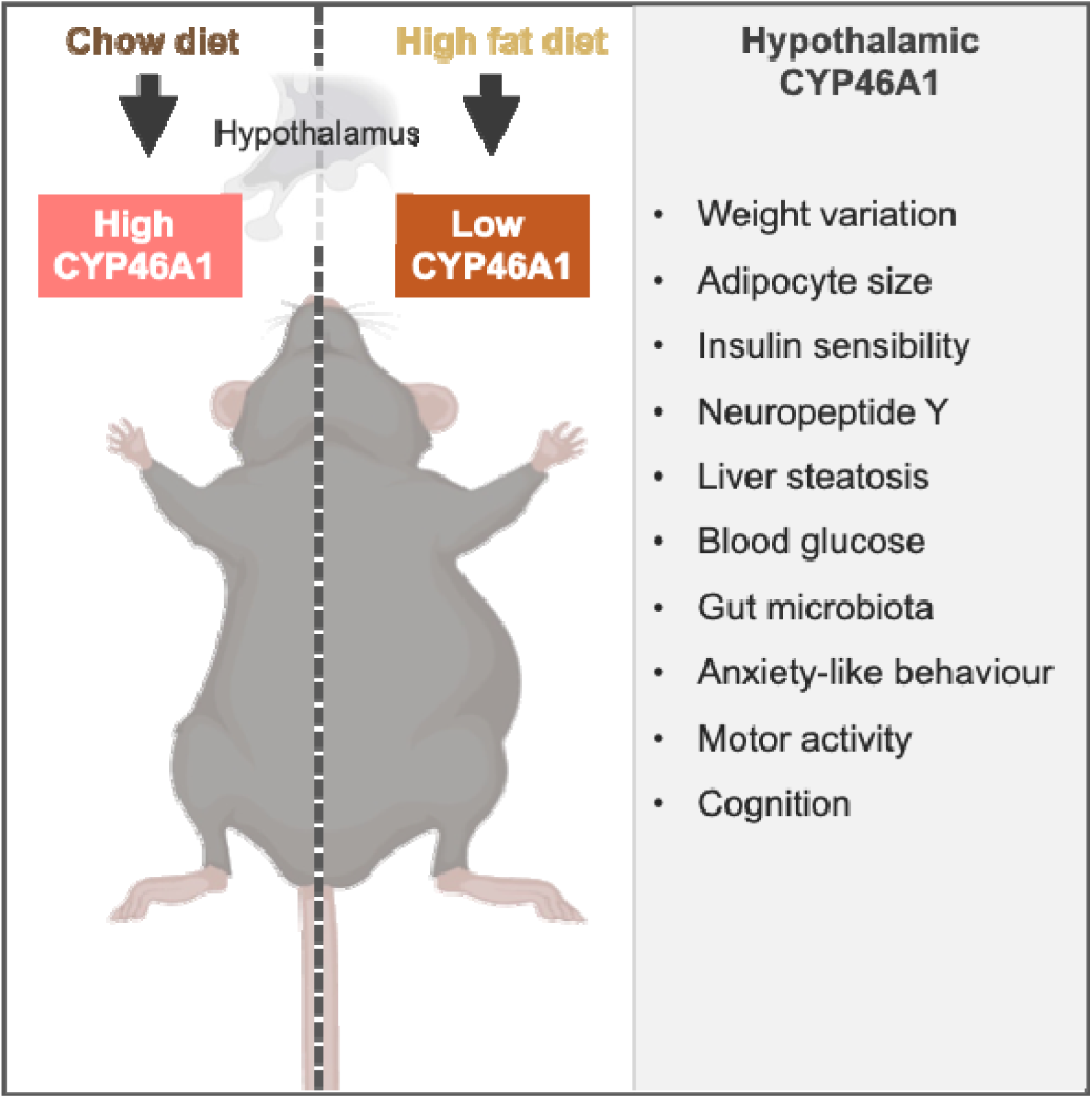
Graphical representation of main findings.

Cholesterol metabolism plays a critical role in brain physiology. About 70%-80% of brain cholesterol is concentrated in myelin sheaths, which are critical for axonal insulation, maintenance of neuronal and efficient synaptic transmission^34^. Cholesterol is also fundamental to preserve structural integrity and fluidity of neuronal cell membranes, enabling proper receptor function and signal transduction. It also contributes to synapse formation and plasticity and serves as a precursor for steroid hormones and bioactive metabolites that support energy production and cellular homeostasis^35,36^. Therefore, the balance between *de novo* cholesterol synthesis and cholesterol efflux is a tightly regulated process in the brain^35^. Cholesterol cannot cross the blood–brain barrier, making brain cholesterol homeostasis highly reliant on local synthesis and turnover, a process in which the enzyme CYP46A1 plays a pivotal role by converting cholesterol into 24S-hydroxycholesterol to enable its elimination. Dysfunction in this balance has been reported in several neurodegenerative disorders (Huntington’s disease, Alzheimer’s disease, Spinocerebellar ataxia and RETT syndrome)^21,36–38^ making CYP46A1, an enzyme that plays a pivotal role in the regulation of brain cholesterol metabolism, a promising therapeutic target. The CYP46A1 enzyme hydroxylates cholesterol into 24-hydroxycholesterol (24-OHC), which can then cross the blood brain barrier for peripheral secretion^39^. We have shown in past studies that viral-mediated knockdown of CYP46A1 leads to an accumulation of cholesterol and a decrease in 24-OHC levels, which are associated with cell death as well as both motor and cognitive impairments^10,21,40^. Therefore, alterations in the levels of 24-OHC and cholesterol can be suggestive of metabolic changes linked to CYP46A1 enzyme activity. Past studies have reported decreased levels of oxysterol in diet-induced and genetic models of obesity, as well as hypothalamic alterations, including inflammation^41^. The ARC of the hypothalamus is a key region involved in the regulation of whole-body energy metabolism and oxysterols levels ^42,43^. The ARC is located adjacent to the median eminence, a circumventricular region characterized by fenestrated capillaries. It is composed of populations of first-order neurons that integrate peripheral signals and, in response, produce and release neuropeptides and neurotransmitters to regulate whole-body energy homeostasis^44^. We hypothesized that diet-induced obesity could, in part, be mediated by modulation of CYP46A1 expression and the downstream reduction of 24-OHC levels and cholesterol accumulation in ARC neurons, thereby contributing to the development of obesity.

HFD-induced obesity has been shown to decrease oxysterols levels in the hypothalamus, particularly 4β-hydroxycholesterol (4β-OHC). Nonetheless, no changes were detected for 24-OHC or CYP46A1 levels in mice that underwent HFD in a past study^41^. In our study, we found that HFD induces a significant reduction of hypothalamic CYP46A1 mRNA compared to animals that undertook chow diet. This apparent discrepancy could be attributed to differences in experimental design, including the use of different mouse strains, which were not specified in the previous study. Indeed, mimicking the HFD-associated decrease in CYP46A1 expression led to increased food and water intake, along with marked body weight gain, suggesting that silencing the *Cyp46a1* gene can recapitulate key effects of an HFD. These effects were further supported by overexpression experiments, which produced opposite phenotypes. CYP46A1 knockdown led to a significant increase in WAT weight, accompanied by adipocyte hypertrophy, further supporting the notion that silencing the *Cyp46a1* gene mimics the metabolic effects of an HFD. The transcription factor PPAR-γ regulates the expression of adipose-specific genes, which play a role in adipogenesis and lipid accumulation within adipocytes^45^. PPAR-γ knockout mice lack adipose tissue due to impaired adipocytes differentiation^46,47^. Beyond its role in adipogenesis, PPAR-γ activation past studies shown that it also contributes to the regulation of insulin sensitivity and is involved in the inhibition of proinflammatory cytokines^48,49^. In this study, we found that CYP46A1 modulates the levels of PPAR-γ protein in WAT. These findings are in accordance with previous reports showing that PPAR-γ deficiency in adipose tissue is associated with increased susceptibility to insulin resistance and lipodystrophy^50,51^. It is tempting to speculate that the variation in PPAR-γ levels may be linked to an inflammatory state in the adipose tissue of these animals. This possibility is supported by the well-established role of PPAR-γ activation in inhibiting the expression of proinflammatory genes in adipose tissue macrophages ^52,53^. Moreover, our results show that shCyp46a1-injected mice exhibited reduced PPAR-γ levels, which correlates with weight gain and adipocyte hypertrophy. This could represent a potential mechanism to counteract fatty acid uptake^50^. It is well established that mice and rats that undergo HFD show reduced hypothalamic NPY mRNA levels^54–58^, possibly as a compensatory mechanism to limit food intake. Our results also replicated this finding in mice that underwent HFD. shCyp46a1-injected mice recapitulated this NPY mRNA decrease, suggesting that HFD decreases CYP46A1 levels, which in turn mediate the reduced expression of NPY in the hypothalamus. Accordingly, CYP46A1 overexpression promotes the opposite phonotype. Overall, these results suggest that hypothalamic CYP46A1 expression regulates key master regulators of the balance between energy consumption and expenditure, which are crucial for maintaining whole-body energy homeostasis.

ARC dysfunction in insulin signaling disrupts whole-body energy homeostasis, leading to impaired glucose homeostasis and ultimately, obesity^34^. Studies have shown that mice undergoing HFD have body weight gain, impaired glucose tolerance and insulin resistance^59–61^, which is consistent with our results. shCyp46a1-injected animals displayed a decrease in glucose tolerance and insulin sensitivity suggesting that the silencing of the *Cyp46a1* gene in ARC, results in dysfunctional insulin signaling. Moreover, CYP46A1 levels in the hypothalamus influence the overall weight of several organs, likely through modulation of lipid content. Adaptive responses in endocrine function occur when the storage capacity of the adipose tissue is exceeded, particularly through increased secretion of adipocytokines and ectopic fat accumulation^62^. This accumulation leads to lipotoxicity, which promotes low-grade inflammation and metabolic dysfunctions, including insulin resistance in several organs^62^. The obesity phenotype observed when decreasing CYP46A1 levels is strongly associated with lipid accumulation in the liver, presenting a risk factor for the development of fatty liver diseases, such as nonalcoholic fatty liver disease^62^. Indeed, we observed an increase in liver weight accompanied by lipid accumulation in hepatocytes, characterized by both macrovesicular and microvesicular steatosis, as well as hepatocytes hypertrophy. Of note, metabolic dysfunction in adipose tissue, liver, and pancreas may also contribute to the decreased insulin sensitivity observed in HFD animals by reducing hypothalamic Cyp46a1 levels.

The gut microbiota is increasingly recognized as a key mediator of obesity and metabolic dysfunction, particularly through bidirectional communication along the gut–brain axis that is sensitive to dietary composition^29^. The results of the present study suggest that hypothalamic Cyp46a1 levels influence gut microbiota composition, particularly *Actinobacteria* and *Tenericutes*, in response to dietary interventions. While no significant differences were observed in Bacteroidetes, Firmicutes, SRB, Betaproteobacteria, and Delta- and Gammaproteobacteria, the Actinobacteria phylum showed a diet-dependent decrease in HFD-fed mice, which was further pronounced in the shCyp46a1 group. This aligns with previous studies reporting a reduction in *Actinobacteria* following HFD exposure^63,64^. However, contradictory findings exist, as some studies associate HFD with an increase in Actinobacteria^65,66^, linking it to body weight gain and pro-inflammatory cytokine levels. The observed variability in *Actinobacteria* responses across studies may indicate the involvement of additional regulatory mechanisms beyond diet alone, potentially including host metabolic and immune factors. *Tenericutes* levels remained unchanged between diet groups, except for a reduction in CYP46A1-overexpressing mice across both diets. This is consistent with prior with reports of increased *Tenericutes (Mollicutes)*^67^ in non-obese mice but contrasts with findings showing HFD-driven reductions in Tenericutes^68,69^.These literature inconsistencies highlight the complexity of diet-microbiome interactions and the potential modulatory role of hypothalamic Cyp46a1. These literature inconsistencies highlight the complexity of diet-microbiome interactions and the potential modulatory role of hypothalamic Cyp46a1.

It has been reported that mice fed a HFD are more immobile and exhibit anxiety-like behavior, such as spending less time in the central zone of the open field^70^. The decrease in total distance traveled and the increase in the time spent immobile suggest a reduction in energy expenditure, possibly due to the increase in body weight. Mice with reduced Cyp46a1 levels spent less time spent in the center, showed increased immobility, and displayed elevated grooming behavior. Conversely, mice overexpressing CYP46A1 exhibited the opposite results, underscoring the importance of cholesterol metabolism in regulating whole-body homeostasis. A study reported that HFD negatively affects hypothalamic function and impairs spatial working memory and increasing anxiety-like behavior in C57BL/6 mice^71^. Our data showed that shCyp46a1-injected mice performed worse in the Y-maze spontaneous alternation test, while CYP46A1-overexpressing mice exhibited improved working memory. These results are in line with a previous study reporting that hippocampal reduction of CYP46A1 levels induced cognitive impairment and hippocampal atrophy^10^ while AAV-mediated overexpression of CYP46A1 improved cognition in AD mice ^37,72^.These independent results suggest that CYP46A1 impacts cognitive performance and behavior through different mechanisms.

In summary, this study places CYP46A1 as a crucial hypothalamic regulator of metabolic and behavioral responses to high-caloric intake. Our findings highlight a novel mechanistic link between hypothalamic cholesterol metabolism and whole-body energy homeostasis. The ability of CYP46A1 to regulate key metabolic pathways suggests that its modulation could represent a potential therapeutic avenue to treat obesity and related metabolic disorders.

## Acknowledgments

This work was supported by the Algarve Biomedical Center Research Institute Funds and by the Molecular Neuroscience and Gene Therapy laboratory funds.

---

**Supplementary Figure 1.**
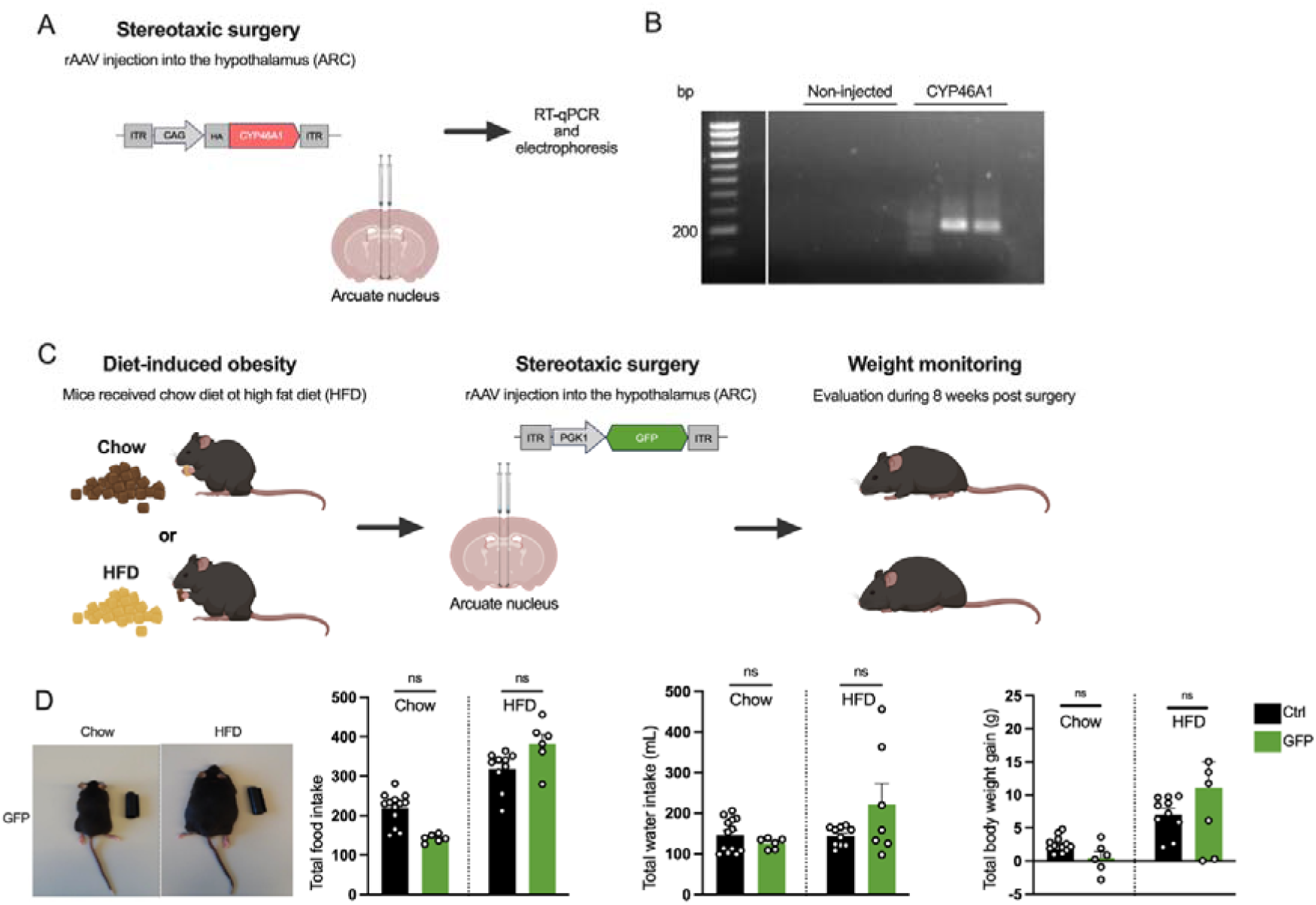
CYP46A1 overexpression validation and control for hypothalamic protein overexpression. **A**) Experimental scheme. Mice were stereotaxically injected with AAV expressing GFP in the arcuate nucleus of the hypothalamus. (**B**) The hypothalamus of mice was microdissected and levels of the human CYP46A1 gene were amplified and visualized in an agarose gel. (**C**) Experimental scheme. Mice underwent chow or high fat diet (HFD) for 4 weeks and were stereotaxically injected with AAVs expressing GFP in the arcuate nucleus of the hypothalamus. mice were evaluated for 8 weeks post-surgery for food or water intake variation and body weight (N=6-13). One-way ANOVA; Fisher’s test. Ns, nonsignificant. PGK1, phosphoglycerate kinase 1.

**Supplementary Figure 2.**
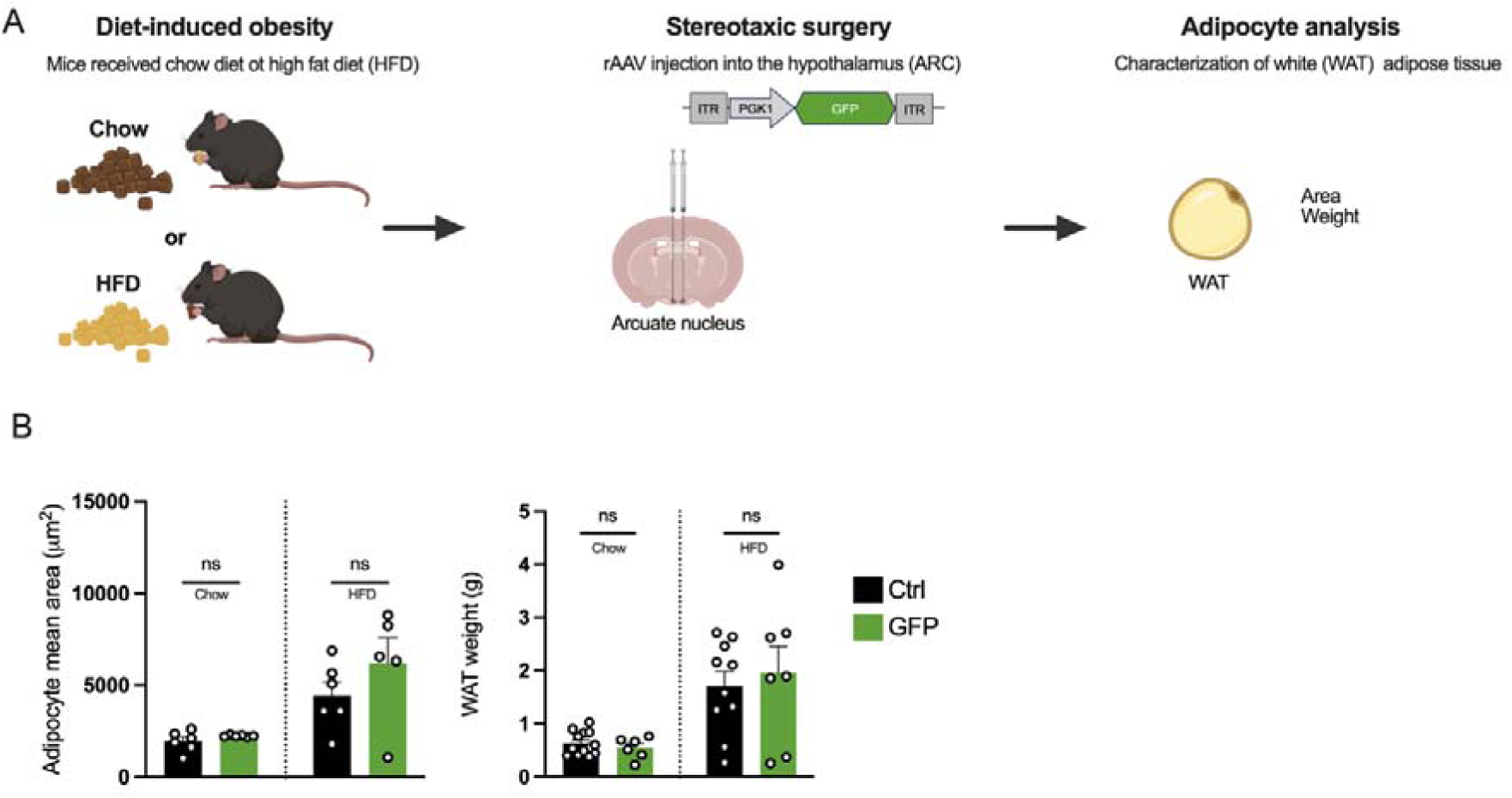
Hypothalamic protein overexpression per se does not affect adipocyte size. (**A**) Experimental scheme. (**B**) Mice underwent chow or high fat diet (HFD) for 4 weeks and were stereotaxically injected with AAVs GFP in the arcuate nucleus of the hypothalamus. White adipose tissue (WAT) was isolated for histological analysis of adipocyte area (N=5-6) and weight (N=6-13). One-way ANOVA; Fisher’s test. Ns, nonsignificant. PGK1, phosphoglycerate kinase 1.

**Supplementary Figure 3.**
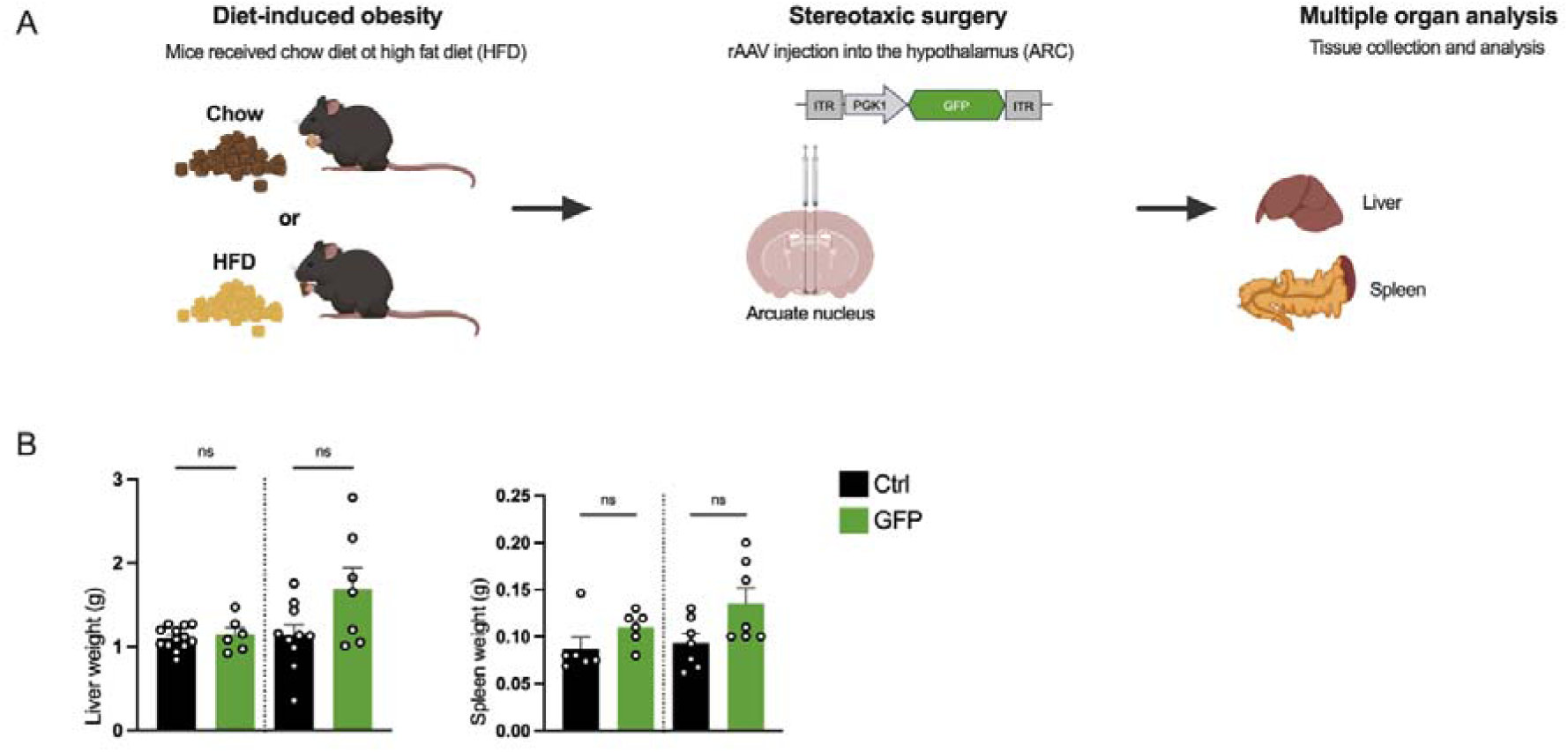
Hypothalamic protein overexpression per se does not affect liver or spleen weight. (**A**) Experimental scheme. (**B**) Mice underwent chow or high fat diet (HFD) for 4 weeks and were stereotaxically injected with AAVs GFP in the arcuate nucleus of the hypothalamus. Liver (6-13). and spleen (6-7) were weighted. One-way ANOVA; Fisher’s test. Ns, nonsignificant. PGK1, phosphoglycerate kinase 1.

**Supplementary Figure 4.**
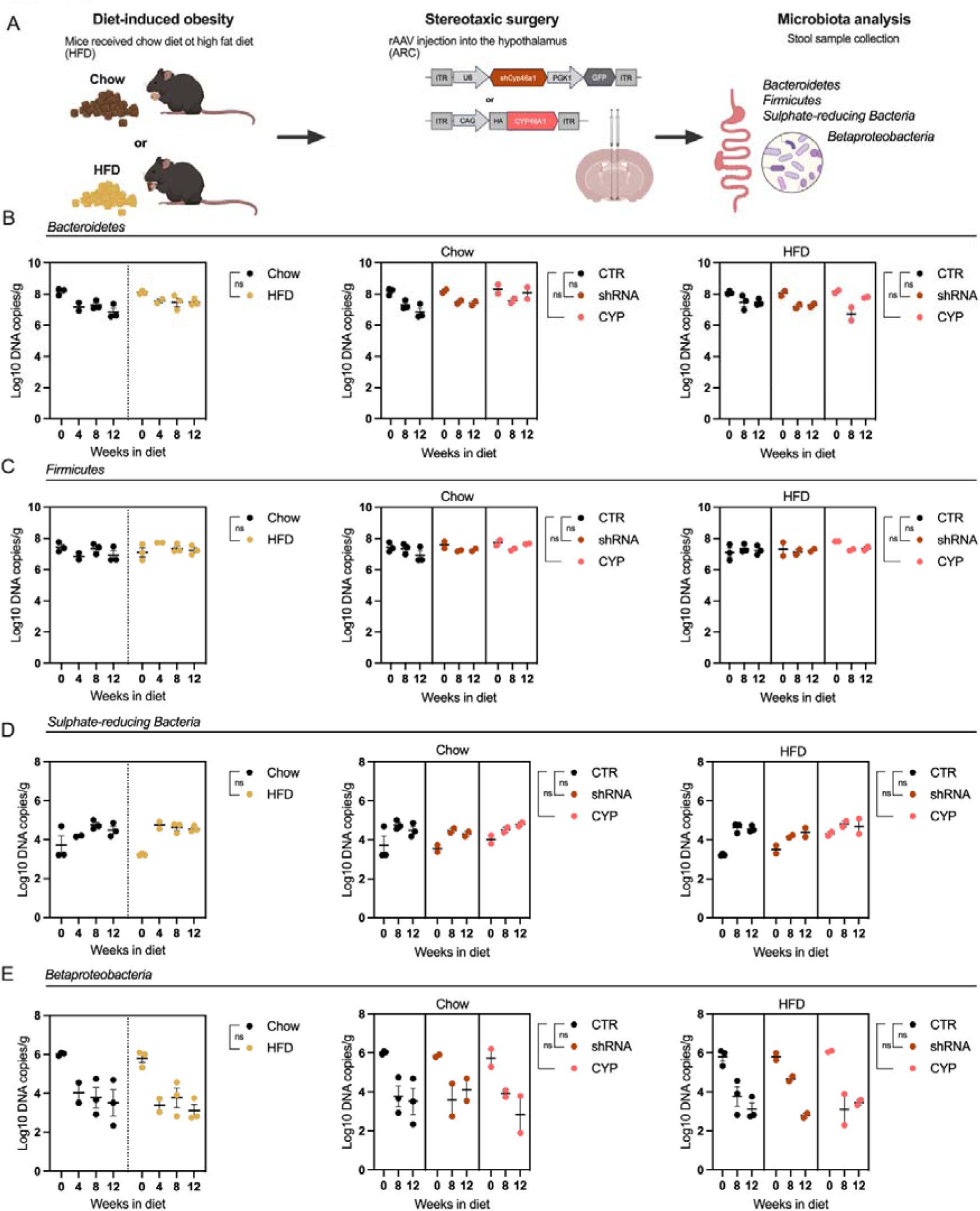
Hypothalamic Cyp46a1 does not alter overall microbiota composition (**A**) Experimental scheme. Mice underwent chow or high fat diet (HFD) for 4 weeks and were stereotaxically injected with AAVs expressing a shCyp46a1 or CYP46A1 in the arcuate nucleus of the hypothalamus. Stool samples from mice were collected and the counts of bacterial taxa by qPCR of 4 bacterial groups as a measure of gut microbiota: *Bacteroidetes, Firmicutes, Sulphate-reducing Bacteria, Betaproteobacteria*. Evaluation by qPCR according to diet (Chow vs HFD, left) (N=11) or CYP46A1 modulation (shCyp46a1 or CYP46A1 overexpression) (N=6-9) of Phyla levels: **(B)** *Bacteroidetes*, **(C)** *Firmicutes*, **(D)** *Sulphate-reducing Bacteria* and **(E)** *Betaproteobacteria*. Nested Student’s t test, Ns, nonsignificant.

